# Cellular organization of visual information processing channels in the mouse visual cortex

**DOI:** 10.1101/441014

**Authors:** Xu Han, Ben Vermaercke, Vincent Bonin

## Abstract

Visual processing and behavior depend on specialized neural representations and information channels that encode distinct visual information and enable distinct computations. Our understanding of the neural substrate, however, remain severely limited by sparse recordings and the restricted range of visual areas and visual stimuli considered. We characterized in the mouse the multidimensional spatiotemporal tuning properties of > 30,000 layer 2/3 pyramidal neurons across seven areas of the cortex. The dataset reveals population specialized for processing of oriented and non-oriented contrast, spatiotemporal frequency, and motion speed. Areal analysis reveals profound functional diversity and specificity as well as highly specific representations of visual processing channels in distinct visual areas. Clustering analysis shows a branching of visual representations along the posterior to anterior axis, and between lateral and dorsal areas. Overall, this dataset provides a cellular-resolution atlas for understanding organizing principles underlying sensory representations across the cortex.

**Summary:** Visual representations and visual channels are the cornerstones of mammalian visual processing and critical for a range of life sustaining behaviors. However, the lack of data sets spanning multiple visual areas preclude unambiguous identification of visual processing streams and the sparse, singular recording data sets obtained thus far are insufficient to reveal the functional diversity of visual areas and to study visual information channels. We characterized the tunings of over 30,000 cortical excitatory neurons from 7 visual areas to a broad array of stimuli and studied their responses in terms of their ability to encode orientation, spatiotemporal contrast and visual motion speed. We found all mouse visual cortical areas convey diverse information but show distinct biases in terms of numbers of neurons tuned to particular spatiotemporal features. Neurons in visual areas differ in their spatiotemporal tuning but also in their relative response to oriented and unoriented contrast. We uncovered a population that preferentially responds to unoriented contrast and shows only weak responses to oriented stimuli. This population is strongly overrepresented in certain areas (V1, LM and LI) and underrepresented in others (AL, RL, AM, and PM). Spatiotemporal tunings are broadly distributed in all visual areas indicating that all areas have access to broad spatiotemporal information. However, individual areas show specific biases. While V1 is heavily biased in favor of low spatial and temporal frequencies, area LM responds more strongly to mid-range frequencies. Areas PM and LI are biased in favor of slowly-varying high-resolution signals. By comparison, anterior areas AL, RL and AM are heavily biased in favor of fast-varying, low to mid spatial frequency signals. Critically, theses biases express themselves in vastly different number of cells tuned to particular features, suggesting differential sampling of visual processing channels across areas. Comparing across areas, we found divergent visual representations between anterior and posterior areas, and between lateral and dorsal areas, suggesting the segregated organization of cortical streams for distinct information processing.

## Introduction

The visual cortex is one of the most elaborate sensory processing network and processing in its multiple areas are important for a range of perceptual, cognitive and motor abilities (Dipoppa et al., 2018; Marques et al., 2018; Minderer et al., 2019; Steinmetz et al., 2019). How distinct visual areas integrate and carry information from distinct visual channels is fundamental to their function and constrains the computations they can perform. However, their functional organization and composition particularly at the cellular level is largely unknown.

Studies of the mouse cortex have the potential to shed significant new light on visual information processing streams and their contributions to perception and behavior. The mouse visual cortex contains over ten higher visual areas (HVAs), which receive direct input from primary visual cortex (V1) and form parallel representations of the visual field (Garrett et al., 2014; Wang and Burkhalter, 2007; Zhuang et al., 2017). HVAs display rich functional response properties and connectivity allowing for a wide range of associations and behaviors (Gamanut et al., 2018; Juavinett and Callaway, 2015; La Chioma et al., 2019; Marshel et al., 2011; Minderer et al., 2019; Olcese et al., 2013; Roth et al., 2012; Smith et al., 2017; Tohmi et al., 2014). HVAs show differences in visual tuning properties including orientation selectivity, spatiotemporal frequency and motion speed preference (Andermann et al., 2011; Marshel et al., 2011; Murakami et al., 2017). Areas PM and AL, for example, show marked preferences for slow and fast varying stimuli and neurons projecting to these areas show functional differences that are biased towards the preferences of target areas (Glickfeld et al., 2013; Han et al., 2018; Kim et al., 2020, 2018). The specificity of visual representations and long-range connections are suggestive of specialized subnetworks involving distinct visual processing channels.

In primates there is anatomical and functional evidence of visual processing channels, namely ventral and dorsal streams, that are specialized in processing distinct visual features at different levels of complexity along the visual hierarchy (Felleman and Van Essen, 1991; Nassi and Callaway, 2009). Whether such distinct visual processing channels exist in the mouse cortex and the extent to which distinct cortical areas partake in these processing channels is yet to be determined. Existing datasets, whether in rodents or in primates although compelling, vastly undersample the cellular functional diversity of visual cortical areas. To study diversity in visual areas and reveal visual information processing channels, it is essential to directly compare response patterns across visual areas and to employ a broad enough stimulus set and large neuronal sampling to allow characterization of a wide range of tuning properties of visual neurons. Such sampling would allow classification of visual neurons into functional cell types and identification of shared cell classes across visual areas to uncover distinct processing channels (Baden et al., 2016; Piscopo et al., 2013; Román Rosón et al., 2019). The vast majority of studies in mouse visual cortical areas used gratings stimuli, which only robustly activate a fraction of the neurons. A recent large-scale study (de Vries et al., 2020) recorded responses to broader sets of stimuli and found different visual areas and cell types show differential stimulus selectivity, but did not address visual channels.

To characterize the functional diversity of mouse visual areas and probe their functional specificity and involvement distinct visual information processing channels, we characterized the multidimensional tuning properties of > 30,000 layer 2/3 pyramidal neurons in 7 mouse visual areas. We found that all mouse visual cortical areas encode diverse albeit distinct visual information. The distinct encoding and functional diversity are seen in overall bias of visual tuning and the sampling of visual features. Individual visual areas encode distinct visual motion speeds and also differentially partake in distinct visual information processing channels. Altogether, this study shows profound diversity and specificity of representations in mouse visual cortical areas and provides a cellular-resolution blueprint for understanding organizing principles underlying visual cortical processing.

## Results

### Characterizing layer 2/3 neurons with cellular imaging and visual noise stimuli

We used large-scale chronic cellular imaging (Figure 1A-C) to probe the visual tuning properties of layer 2/3 cortical excitatory neurons in 7 mouse visual areas (V1; LM: lateromedial; AL: anterolateral; RL: rostrolateral; AM: anteromedial; PM: posteromedial; LI: laterointermediate). We focused on tuning for orientation and spatiotemporal features that are characteristic of visual information processing channels in the mammalian visual system. In the first part of this paper, we examine, for each of the studied visual areas, both the biases in neuronal tuning and the neural representation of the visual feature space. In the second part, we examine the implications of cortical neurons’ visual tuning for encoding of visual motion speed and for the organization of visual information processing channels in the mouse cortex.

**Figure 1.**
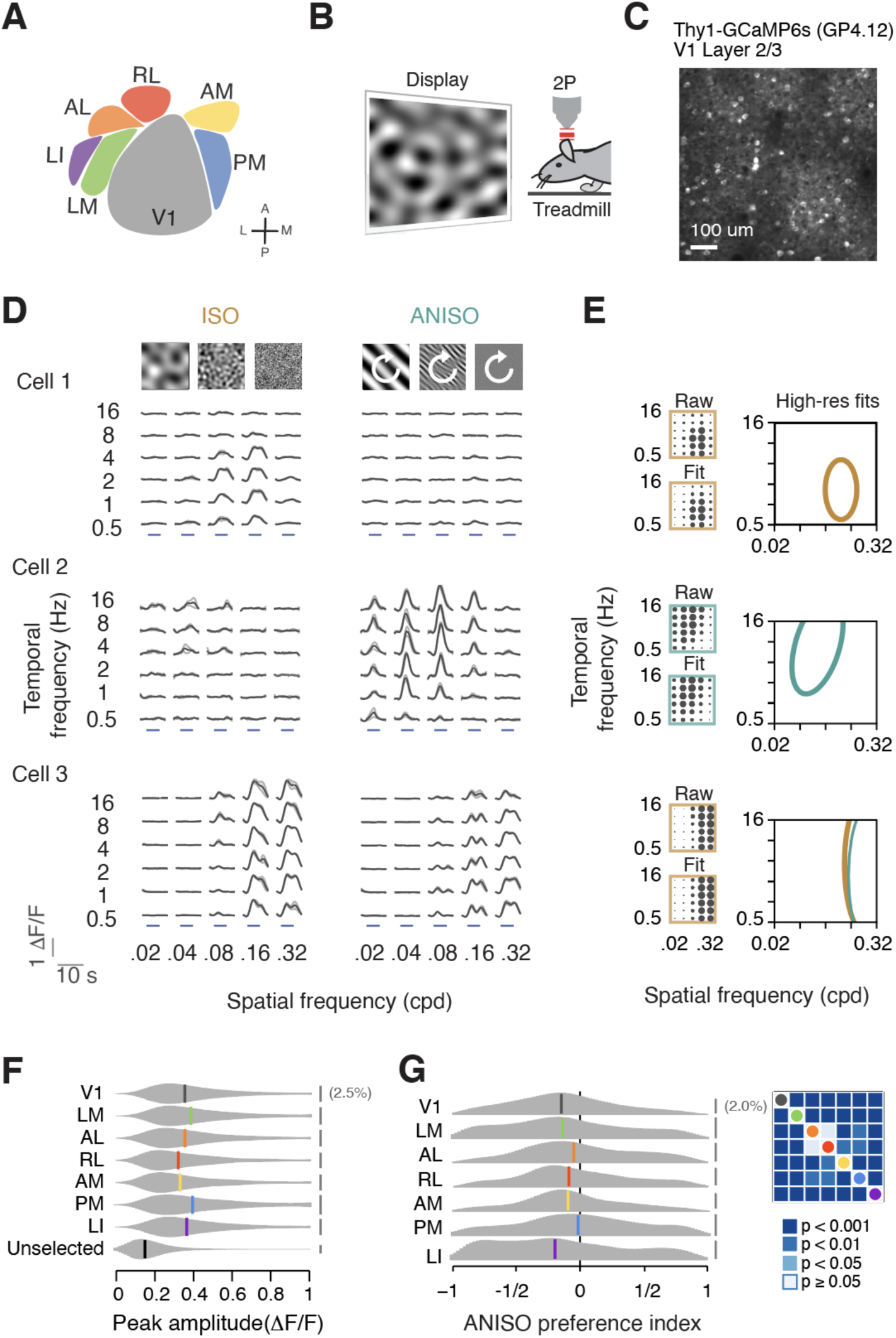
Spectral noise stimuli elicited rich responses in multiple visual cortical areas. (A) Cartoon showing V1 and six higher visual areas investigated in this study. (B) Experimental setup for 2-photon calcium imaging on awake mice. Mice were placed on a treadmill while cellular responses to spectral noise stimuli were recorded using 2-photon calcium imaging. (C) Example field of view of the V1 L2/3 pyramidal neurons of Thy-GCaMP6s (GP4.12) mice. Scale bar: 100um. (D) Example cells showing distinct responses to ISO and ANISO stimuli (isotropic vs anisotropic noise stimuli; left and right icons). ISO and ANISO stimuli have varying spatiotemporal frequencies, and non-oriented and oriented textures, respectively. See methods and figure supplement 1 for a detailed description. Gray traces show the median stimulus-triggered responses across trials. The median absolute deviations across trials are shown as light gray shadows. Cell 1 and 2 selectively responded to ISO and ANISO stimuli respectively, while cell 3 showed similar responses to both stimuli. Blue bars indicate 4-sec stimulus epochs. ΔF/F: ratio of fluorescent changes. (E) Model fitting of spatiotemporal frequency tunings. Neurons’ responses to the preferred stimuli are used for fitting. Yellow: ISO response; green: ANISO responses. The calcium responses during the stimulus epoch are averaged to generate the raw response maps (upper left), which are then fitted to two-dimensional Gaussian models (bottom left). The size of the dot indicates the normalized response amplitude. A high-resolution fit is generated by oversampling (right panel). Contours indicate the region of frequency eliciting responses greater than 50% of the maximal amplitude. (F) Distribution of peak responses of reliable cells and unselected cells. Distributions are smoothed with a Gaussian kernel (bandwidth=0.05). Color bars indicate median values. Gray scale bars indicate 2.5% of cells of each distribution. (G) Distribution of ANISO preference index of each area. Gaussian kernel bandwidth=0.05. Gray scale bars indicate 2% of cells for each distribution. Pairwise statistical comparisons were shown in the right panel (KS test with Bonferroni correction). See Table 1 for sample sizes. The following figure supplements are available for figure 1: Figure supplement 1: Retinotopic mapping and spectral noise stimuli. Figure supplement 2: Population statistics of response to ISO and ANISO stimuli.

To characterize neuronal tunings for visual features in mouse visual cortical areas, we used large-field stochastic visual stimuli designed to activate diverse orientation and spatiotemporal frequency channels. The stimuli were made of spectrally-filtered random noise with filters of specific center spatial and temporal frequencies, center orientation, and orientation bandwidth (Figure supplement 1D; Materials and Methods), resulting in dynamic textures that varied in orientation, spatiotemporal frequency and spatial elongation (anisotropy). We catalogued in separate imaging sessions responses to isotropic (non-oriented) textures and anisotropic (oriented) textures of different spatiotemporal frequencies (Figure 1D). To characterize functionally-diverse populations and probe visual processing channels, we tested 30 different combinations of spatial and temporal frequencies (spatial frequencies: 0.02, 0.04, 0.08, 0.16, 0.32 cpd; temporal frequencies: 0.5, 1, 2, 4, 8, 16 Hz; data sets 1 and 2). The power spectrum of the isotropic textures (Figure 1D, left) is evenly distributed across spatial orientations and therefore well suited to probe neurons tuned to non-oriented features. The anisotropic stimuli (Figure 1D, right) had power restricted to a narrow band of orientations (5 deg, full width at half maximum, FWHM) and are well suited to probe the properties of orientation tuned neurons. To cover a full range of orientations, the stimulus center orientation was varied slowly (45 deg/s) over stimulation epochs. In a separate experiment, we recorded responses to stimuli of different elongation and spatial frequencies (data set 3). The visual stimuli were presented on a visual display positioned in the right visual field (azimuth 0∼100 deg; elevation -30∼50 deg) in 4-second epochs interleaved with 4-second epochs of gray screen of the same mean luminance. The stimuli were presented four times in randomized order with stimuli in repeated trials obtained using distinct pseudo-random generator seeds. Data sets were registered to common reference to match somatic ROIs across experiments.

**Table 1.** Population for data analysis. Table summarizing the number of mice and cells imagined for each area, responsive cells (response amplitude > 3sd of baseline ΔF/F0 for over 1 sec for at least one stimulus condition), cells showing reliable responses (75^th^ percentile of cross-trial correlation coefficient > 0.3) to individual and combinations of datasets, cells with good fits of spatiotemporal responses.

| Area | # Mice | # Cells | Responsive<br>(% all cells)<br>Dataset 1-3 | Reliable responses (% all cells) |  |  |  |  | SFTF good fits |  |  |
| --- | --- | --- | --- | --- | --- | --- | --- | --- | --- | --- | --- |
|  |  |  |  | Dataset 1<br>(ISO) | Dataset 2<br>(ANISO) | Dataset 3<br>(Elongation) | Dataset<br>1&2 | Dataset<br>1-3 | Dataset 1 | Dataset 2 | Dataset<br>1&2 |
| V1 | 7 | 15658 | 12515(80) | 5736(37) | 5476(35) | 7309(47) | 7498(48) | 9447(60) | 5421 | 5031 | 7126 |
| LM | 8 | 22795 | 13516(59) | 6069(27) | 5924(26) | 6939(30) | 8395(37) | 9472(42) | 5661 | 5222 | 7895 |
| AL | 6 | 9694 | 4936(51) | 1821(19) | 2308(24) | 2023(21) | 2849(29) | 3158(33) | 1732 | 2111 | 2705 |
| RL | 6 | 6792 | 4261(63) | 1548(23) | 1812(27) | 1985(29) | 2360(35) | 2800(41) | 1471 | 1693 | 2252 |
| AM | 6 | 7399 | 3625(49) | 1197(16) | 1353(18) | 1574(21) | 1755(24) | 2105(28) | 1178 | 1312 | 1724 |
| PM | 6 | 9275 | 3623(39) | 940(10) | 1100(12) | 1200(13) | 1505(16) | 1802(19) | 927 | 1079 | 1482 |
| LI | 7 | 7317 | 3693(50) | 1422(19) | 1113(15) | 1516(21) | 1905(26) | 2269(31) | 1320 | 1027 | 1802 |
| **Total** | **10** | **78930** | **46169(58)** | **18733(24)** | **19086(24)** | **22546(29)** | **26267(33)** | **31053(39)** | **17710** | **17475** | **24986** |

To target cortical excitatory neurons, we employed transgenic mice (Thy1-GCaMP6s GP4.12, Dana et al., 2014) that broadly express the calcium indicator GCaMP6s in a large subset of pyramidal neurons (Figure 1C). As anesthesia profoundly alters visual response properties (Lien and Scanziani, 2013; Suzuki and Larkum, 2020), recordings were made in awake animals. Mice were head-fixed comfortably on a treadmill, yielding recordings in quiet wakefulness allowing for occasional locomotor behavior. Locomotion epochs were too infrequent to be examined. All data were included irrespective of the animal’s locomotion state. The cellular imaging was targeted to the retinotopic locations in the mouse cortex that corresponded to the visual display center. To identify the borders of visual areas at high spatial resolution, we performed widefield and cellular calcium imaging of responses to retinotopic stimuli (Figure supplement 1A-C). V1 and surrounding higher visual areas were assigned using the reversals in the retinotopic representations (Wang and Burkhalter, 2007; Zhuang et al., 2017). Borders between areas were further refined by manually inspecting cellular responses to retinotopic stimuli in 2-photon images. 2-photon imaging was focused 100–300 μm below the pia to target cortical pyramidal neurons within layer 2/3.

### Specialized populations for processing of oriented and non-oriented contrast

Out of 78,930 identified layer 2/3 neurons (n = 10 mice), 58% showed a stimulus-evoke response to at least one of the visual stimuli tested (46169/78930 cells, > 3sd of the baseline ΔF/F, see Materials and Methods, data sets 1–3). To characterize the neurons’ visual tunings, we identified neurons showing strong, reliable responses to the visual stimuli (figure supplement 2A-B). We correlated the raw ΔF/F activity time courses in visual stimulation epochs across repeated trials and selected the cells showing pronounced trial-to-trial correlations (75th percentile of correlation coefficients > 0.3, see Methods). This criterion served to identify reliably responding cells and exclude weak responders from which clear visual tunings could not be extracted (Figure 1F). Across visual areas, about a third of the probed neurons showed visual responses that fulfilled this criterion (31053/78930 cells, data sets 1–3) yielding samples of 1800 to 9400 functionally-characterized neurons for each visual area (Table 1).

The responses were highly tuned to specific ranges of spatiotemporal frequencies, indicative of visual processing channels (Figure 1D, traces). To characterize these tunings, we fitted 2-dimensional Gaussian functions (Figure 1E) to individual neurons’ trial-averaged amplitudes of stimulus-evoked responses and extracted response descriptors including peak response magnitude, spatial and temporal bandwidths, and preferred and cutoff spatiotemporal frequencies. These function fits captured the magnitudes of responses as well their dependence on spatiotemporal frequency (Figure 1E, left; Figure supplement 2C). For neurons that responded to both isotropic and anisotropic stimuli, spatiotemporal tuning estimates were highly correlated (Figure 1E, bottom; Figure supplement 2D), therefore, we used the fits from the data set with highest peak responses (Figure 1E, thick lines). In total we good fits were obtained for a total of 24986 neurons across areas (normalized residual of fits < 0.1, Table 1).

In addition to spatiotemporal tuning, the neurons also show a variety of responses for isotropic and anisotropic stimuli (Figure 1D). While some cells show similarly strong responses to both types of stimuli (Figure 1D, bottom), many have clear preferences, responding strongly to one stimulus type and weakly to the other (Figure 1D, middle vs. top). A fraction of cells shows a clear preference for anisotropic stimuli (Figure 1D, middle). A preference for anisotropic stimuli is indicative of pronounced sensitivity to oriented edges or orientation tuning, whereas a lack of this preference is indicative of a lack of orientation tuning. Surprisingly, a separate population of visual cortical neurons shows a strong preference for isotropic stimuli, which may reflect tuning for a non-oriented feature. To quantify the neurons’ preference for isotropic and anisotropic stimuli, we extracted peak response amplitudes from model fits, and computed an anisotropy preference index (API), which we defined as the ratio of the difference in peak responses to the stimuli over their sum (Figure 1G). The index has a value of -1 for perfectly isotropic preferring neurons, 1 for perfectly anisotropic preferring neurons and 0 for neurons without preference.

The neurons show highly diverse preferences for isotropic and anisotropic stimuli (Figure 1G -- Figure supplement 2E). Across the population, 32% of the cells showed a clear preference for isotropic stimuli (7928/24986 cells; API < -1/3), whereas 20% showed a clear preference for anisotropic stimuli (5045/24986 cells; API > 1/3). The remaining 48% showed only weak preferences (12013/24986 cells). Importantly, anisotropic and isotropic preferring cells are observed across all visual areas, however, there were specific biases (Figure 1G). Areas V1, LM and LI have a pronounced bias for isotropic stimuli. In contrast, areas AL, RL, AM and PM show less prevalence of isotropic-preferring cells and an enrichment of cells without preference.

Thus, layer 2/3 visual cortical populations show pronounced preferences for isotropic and anisotropic stimuli. This may reflect specific visual channels specialized for oriented and non-oriented contrast.

### Specializations for oriented and non-oriented contrast in dorsal and lateral areas

To examine these visual processing channels in further detail, we characterized, in a separate dataset (22546 cells showing reliable responses, 10 mice), the responses to isotropic stimuli and 3 different levels of anisotropy (orientation bandwidth: infinite, 40, 10, 5 deg FWHM; Figure 2A). We sought to determine whether responses to stimulus elongation form a continuum or a dichotomy. We sampled a range of spatial frequencies (0.04, 0.08, 0.16, 0.32 cpd, 2 Hz), and examined the amplitudes of responses as a function of elongation (Figure 2B).

**Figure 2.**
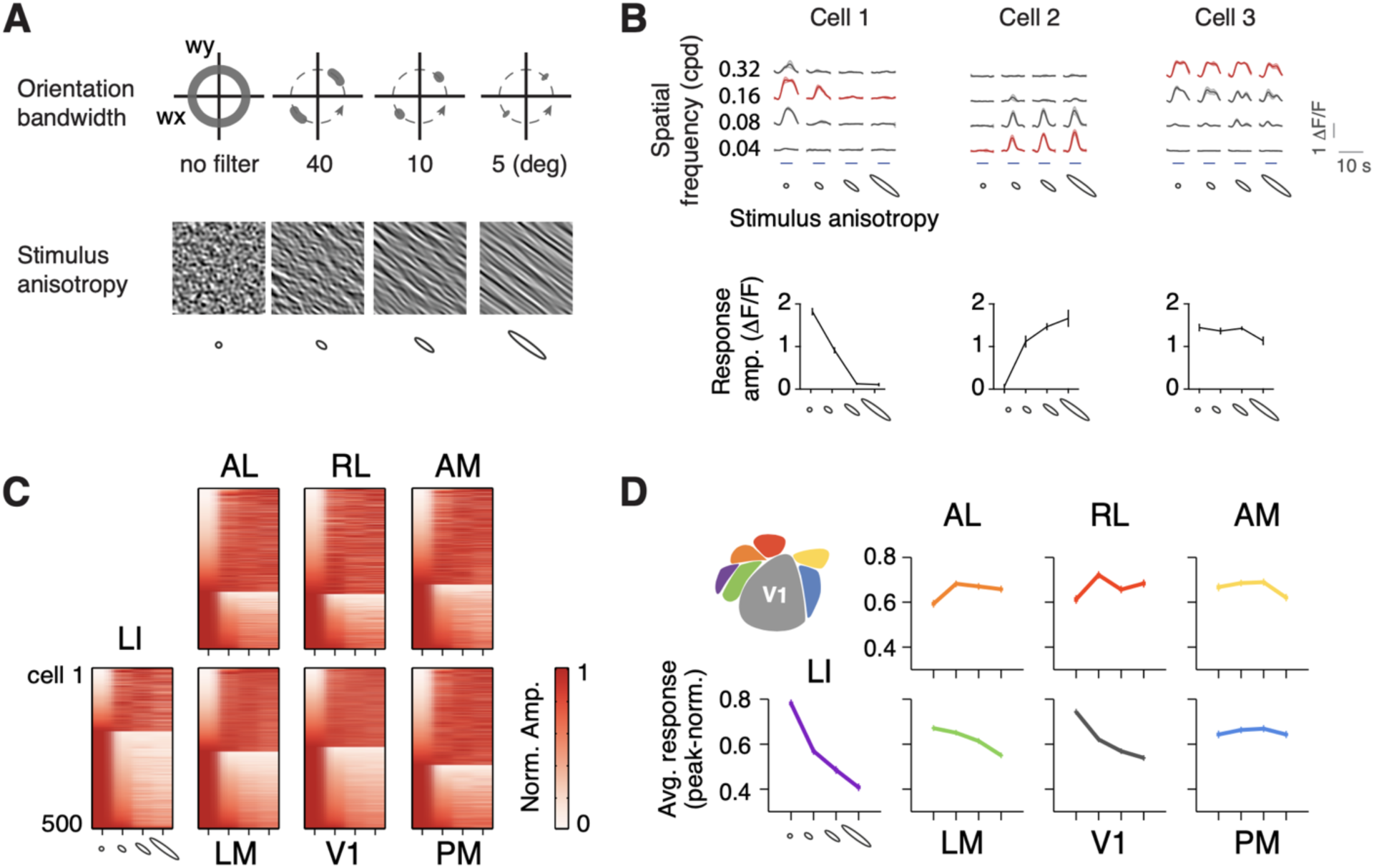
Distinct preferences for oriented and non-oriented stimuli in dorsal and lateral areas. (A) Example noise stimuli showing increasing stimulus anisotropies and texture elongation corresponding to increasingly narrow orientation bandwidths (B) Example cells showing distinct responses to combinations of spatial frequencies and stimulus anisotropies (upper panels). These cells correspond to the example cells in Figure 1D, showing matching response preferences for ISO and ANISO stimuli. Tuning curves for the stimulus anisotropy are measured at the preferred spatial frequencies (red curves) and are shown in the lower panels. (C) Population tuning curves of randomly selected 500 cells from each area, sorted by peak-normalized response amplitude. (D) Average tuning curves for stimulus anisotropy across population of each area. Mean ± sem. Inset: scheme of cortical areas. See Table 1 for sample sizes.

We found across the population a dichotomy of responses to spatial elongation (Figure 2C). Anisotropic preferring cells did not respond to isotropic stimuli (Figure 2B, middle, 2C) and show nearly invariant responses with elongation. Isotropic preferring neurons (Figure 2B, left and right, 2C) instead show a steep fall-off of the response amplitude with elongation. We observed in the population little evidence of tuning to specific length scales. Comparing population response patterns across visual areas (Figure 2C), we found that areas V1, LM, and LI show the largest fractions of cells tuned to isotropic stimuli, whereas areas PM, AL, RL and AM show an enrichment of orientation tuned neurons. The striking differences in the tuning of these areas was also seen in the average tuning curves (Figure 2D).

Taken together the results indicate that visual cortex contains populations specialized for processing of oriented and non-oriented contrast and that these populations are non-uniformly distributed across visual areas.

### Diversity and specificity of spatiotemporal visual channels in visual areas

We next examined the diversity and specificity of spatiotemporal channels in mouse visual areas. Previous studies reported biases in spatiotemporal tunings across visual areas, particularly in areas AL and PM which show biases in favor of fast and slow stimuli respectively (Andermann et al., 2011; Marshel et al., 2011; Murakami et al., 2017). These biases could reflect specific sampling of specific spatiotemporal channels. To examine in detail the diversity and specificity of spatiotemporal tunings in visual areas, lower and higher, we examined population responses to spatiotemporal stimuli, including the numbers of neurons activated as a function of spatiotemporal frequency, and the distributions of spatiotemporal tuning types (lowpass, bandpass, or high-pass), preferred frequencies and tuning bandwidths.

We observed a profound diversity of spatiotemporal channels in mouse visual areas. Each area shows highly diverse tunings, with each neuron individually responding to distinct ranges of spatiotemporal frequencies, but as a population, covering broad swaths of the spatiotemporal spectrum (Figure 3A). This diversity in neuronal tunings is particularly evident in the broad distributions of peak and cutoff spatiotemporal frequencies (Figures 3D; Figure 4A,C; Figure supplement 4A,B), but is also seen in the spatiotemporal tuning types (Figures 3C) and tuning bandwidths (Figure supplement 4C). Critically, the diversity was observed across all of the visual areas, not only seen in lower visual areas V1 and LM, but also in more specialized areas (e.g. AL and PM).

**Figure 3.**
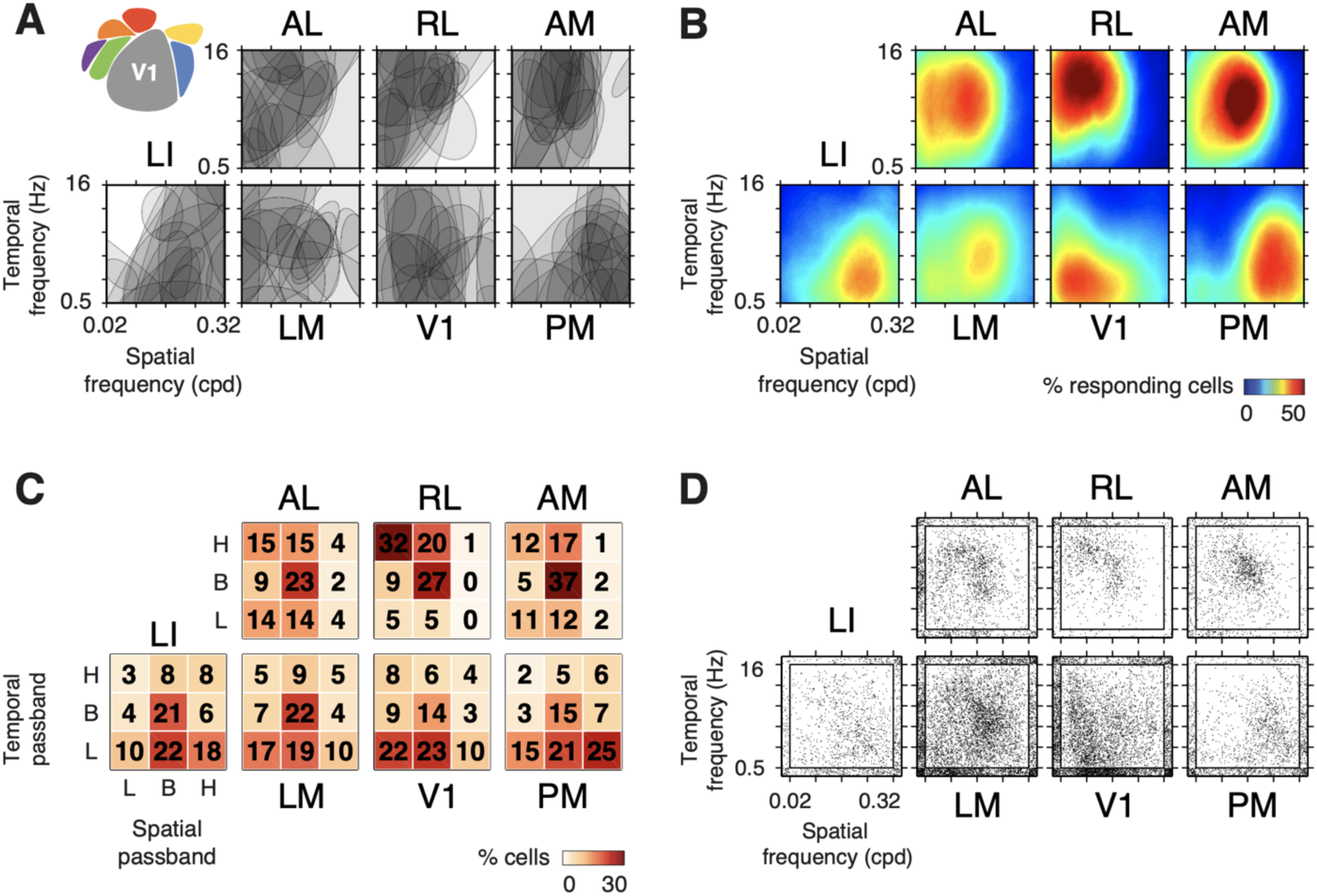
Diversity and specificity of spatiotemporal tunings in visual areas. (A) Superimposed surfaces showing diverse spatiotemporal tunings of 20 randomly selected cells. Inset: scheme of cortical areas. (B) Proportion of neurons responding to certain spatiotemporal frequencies. (C) Proportions of spatiotemporal passband types. H: highpass. B: bandpass. L: lowpass. (D) Scatter plots showing the distribution of preferred frequencies of the populations across areas. The inner box delineates the range of frequency explored in this study, dots at the boarders are scattered outward to visualize the density. See Table 1 for sample sizes.

**Figure 4:**
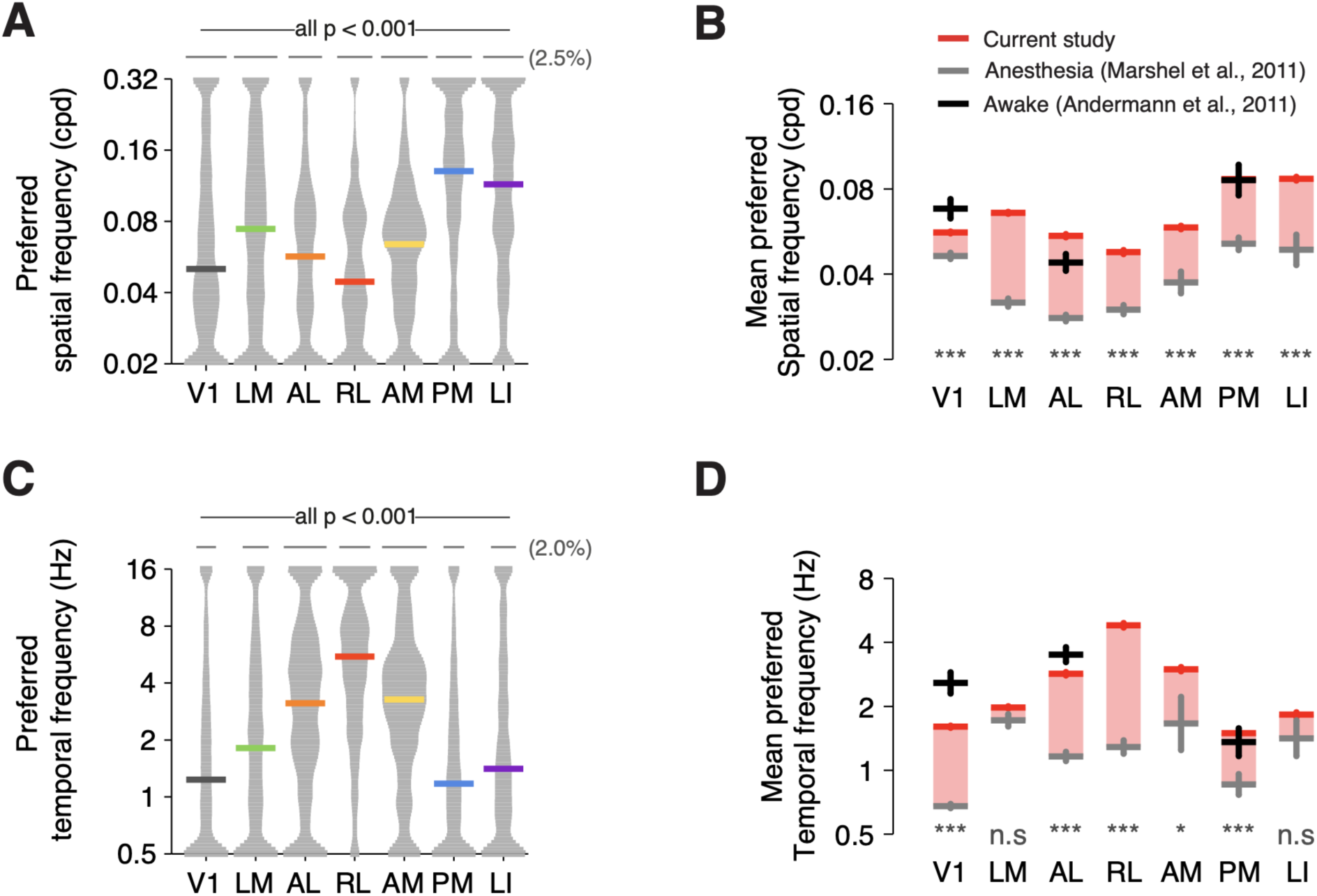
Neurons encoded higher spatial acuity and faster stimuli in wakefulness than in anesthesia. (A) Distributions of preferred spatial frequencies across areas. The width of each column is normalized to the maxima of the distribution. Distributions are smoothed with a Gaussian kernel (bandwidth=0.1). The median values are shown as colored bars. Gray scale bars indicate 2.5% of cells of each distribution. (B) Comparison of geometric means of neurons’ preferred spatial frequencies acquired in this study (red bars) and previous studies in anesthetized (gray bars) and awake mice (black bars). Statistical differences between the current datasets and the ‘anesthesia’ dataset are shown at the bottom row. KS tests with Bonferroni correction. P<0.001, ***; P<0.01, **; P<0.05, *; p ≥ 0.05, n.s. (C-D) Temporal frequency analysis, same as (A-B). See Table 1 for sample sizes. The following figure supplement is available for figure 4: Figure supplement 3: Spatiotemporal tuning parameters.

Although each area is comprised of neurons with diverse spatiotemporal preferences, the biased distributions of these neurons among visual areas give rise to striking functional specificity such that each area displays a unique set of activation patterns in response to spatiotemporal stimuli (Figure 3).

First, V1 and LM, show profound specializations for processing low and mid-range spatio-temporal frequencies, respectively. V1 and LM are differentially activated by low and mid-range frequencies (Figure 3B, **V1** vs **LM**) and differ strikingly in the numbers of spatiotemporally lowpass and bandpass cells (Figure 3C, **V1** vs **LM**) and in their distributions of peak spatiotemporal frequencies (Figure 3D, **V1** vs **LM**). Second, areas PM and LI which are located on opposite sides of V1 share a pronounced preference for low temporal (0.5–4Hz), high spatial frequencies (0.08– 0.32 cpd), which links them to processing of high spatial resolution visual signals. This is seen in the population activation pattern, and the distributions of passband properties and peak frequencies (Figure 3B-D, **PM** and **LI**).

By comparison, areas AL, RL and AM, which are located anterior to V1, share a striking preference for low resolution, fast varying visual signals. These areas show weak activation at low temporal (1 Hz and below) and high spatial frequencies (0.08 cpd and above) (Figure 3B, **AL, RL** and **AM**) and broadly distributed peak frequencies with a mode around 0.08 cpd and 4 Hz (Figure 3D, **AL, RL** and **AM**). They also show a preponderance of temporally bandpass and high-pass neurons and a sparsity of spatially high-pass cells (Figure 3C, **AL, RL** and **AM**). Moreover, anterior visual areas have their own distinguishing features (Figure 3B-D, **AL, RL** and **AM**). Area RL shows an excess of spatially-lowpass and temporally-highpass cells and the strongest activation to stimuli with high temporal frequencies and low spatial frequencies. Area AM, in comparison, shows tunings shifted towards mid-range spatial frequencies and weaker responses to very high temporal frequencies. Area AL comprises a higher diversity of tunings without clear biases seen in areas RL and AM.

These differences between spatial and temporal properties of distinct visual areas of the mouse cortex are robust and surprising. Each area shows a highly distinct distribution of peak spatial and temporal frequencies (Figure 4A,C). Although the preferences measured with ISO and ANISO stimuli are on a par with a study of response properties measured with gratings in V1, AL and PM in awake animals (Figure 4B,D, red vs. black)(Andermann et al., 2011), they differ strikingly from properties measured in anesthetized mice with discrepancies that can exceed one octave (Figure 4B,D, red vs. gray) (Marshel et al., 2011).

These results indicate that spatiotemporal tuning properties of layer 2/3 cortical pyramidal neurons in each mouse visual area show high level of diversity, suggesting distributed processing with a wide range of visual information shared across areas. At the same time unique distributions of tuning parameters in each area creates functional specialization, allowing differential involvement of each area in spatiotemporal information processing. The specialization of HVAs suggests two separate branches of spatiotemporal representations where anterior areas (AL, RL, AM) are involved in processing of fast-moving large objects while lateral areas (LI and PM) are specialized in encoding fine details of still or slowly moving scenes.

### Higher visual areas encode complementary motion speed representations

The distinct spatiotemporal tunings observed in different visual areas suggest that they might encode distinct visual motion speeds. A previous study (Andermann et al., 2011) reported an abundance of speed-tuned cells in area PM, which is biased towards low temporal and high spatial frequencies (Figure 3B). To identify similar biases in our dataset that spans multiple visual areas, we computed a spatiotemporal tuning index ξ extracted from the spatiotemporal tuning fits to the responses to isotropic noise stimuli (Materials and methods). This parameter describes the dependency between neurons’ spatial and temporal tunings, which expresses as the slant of spatiotemporal response profiles (Figure 5A). Neurons with ξ ≥ 0.5 are speed tuned because they show consistent speed tuning curves across broad ranges of spatial frequencies (Figure supplement 4A,B). We examined the distributions of speed tuning indices across visual areas and examined responses of speed-tuned cells to stimuli of different speeds.

**Figure 5.**
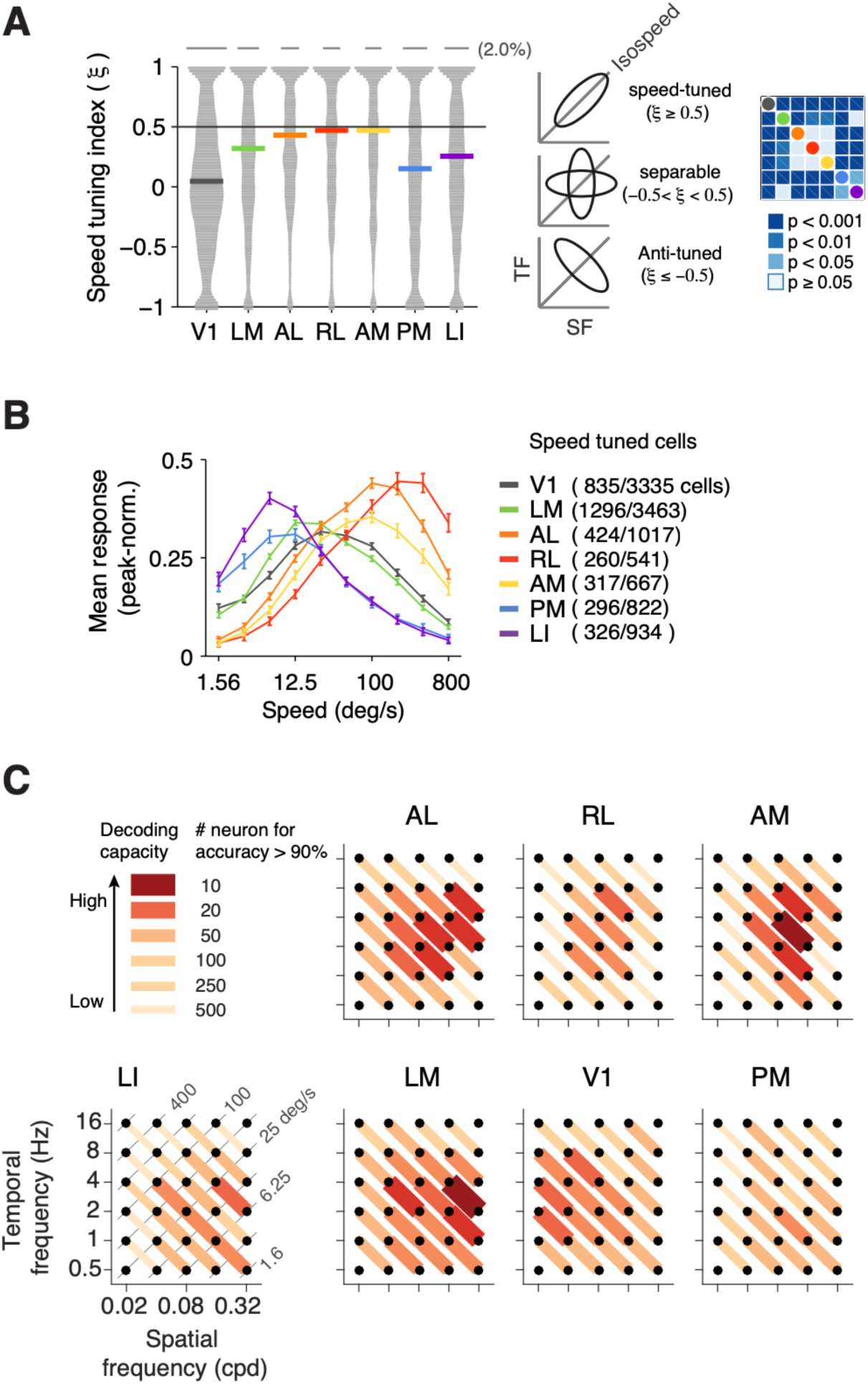
Complementary representation of visual motion speed in higher visual areas. (A) Distribution of speed tuning index ξ of each area. Colored bars show median values. Neurons above the threshold (ξ ≥0.5) are deemed speed-tuned, otherwise untuned. The right panel shows statistical differences of the distributions across areas (KS tests with Bonferroni correction). (B) Population average speed tuning curves of each areas. Each line is the average of peak-normalized speed tuning curves of speed-tuned cells within each area. Error bars: standard error of the mean. (C) Enhanced speed discriminability in higher visual areas. Discrimination tasks were performed between cross-speed pairs of isotropic stimuli, which have 4-fold difference in speed. The discriminability is estimated as the number of cells required to reach 90% classification accuracy (red lines). Higher visual areas (except PM) show regional increases in the discrimination ability comparing to V1. The following figure supplement is available for figure 5: Figure supplement 4: Speed tuning analysis.

We found a striking specialization for visual motion speed in mouse visual areas (Figure 5A). Area V1 shows highly diverse tunings but only a small proportion of speed-tuned cells (25%, 835/3335 cells). Relative to V1, anterior visual areas AL, RL, and AM show a clear enrichment of speed-tuned cells (42∼48%), whereas areas LM, PM and LI show a less profound enrichment (35∼37%). Differences across visual areas are highly significant (Figure 5A, right).

Moreover, speed-tuned neurons encode distinct ranges of speed in different areas (Figure 5B, t-tests). V1 and LM are broadly tuned for the intermediate speeds (peak at 12.5 ∼ 25 deg/s). Areas AL and AM primarily respond to fast-moving stimuli (peak at ∼100 deg/s); RL neurons selectively respond to extremely fast stimuli (peak at 200∼400 deg/s). In contrast, areas PM and LI mainly respond to slow motion (peak at 6.25 deg/s). Therefore, individual higher visual areas are specialized for specific regimes of motion speed and they collectively establish a broad representation of speed that might be critical for a wide range of visual behaviors.

### Enhanced speed discrimination capacity in higher visual areas

To assess whether the visual motion tuning properties endow visual areas with distinct encoding abilities, we compared how well the stimulus categories (spatiotemporal frequencies or speed) can be decoded from the activities of neuronal populations. We trained SVM (support vector machines) classifiers to discriminate stimulus pairs using the responses to ISO stimuli from random cell samples. All recording trials were randomly divided into two halves that were respectively used for training and testing the classifiers (Methods). The discrimination accuracy for each stimulus pair was measured with different population sizes, from which the population size required to reach 90% accuracy was interpolated and used as a proxy of discrimination power of each visual area in discriminating stimulus pairs. To measure the discriminability of speed, we tested how well the classifiers discriminate cross-speed stimulus pairs, which are neighboring stimulus pairs lying orthogonally to the iso-speed lines. The cross-speed pairs have small differences in the spatial (1 octave) and temporal frequency (1 octave), and a modest (4-fold) difference in speed (Figure 5C). Switching the combination of the spatial and temporal frequencies of cross-speed pairs yield iso-speed pairs (Figure supplement 4C), which also have small differences in the spatiotemporal frequency but identical visual motion speed. The difference between the discriminability of cross-speed and iso-speed pairs isolate speed discriminability, which is intermingled with the discriminability of spatiotemporal frequencies.

Higher visual areas showed better speed discrimination power than V1 in distinct frequency domains (Figure 5C). V1 showed relatively uniform ability to discriminate across cross-speed pairs, with a slight increase at lower spatial frequencies. Area LM showed better discrimination at slow to intermediated speed ranges (6.25 – 200 deg/s). Area AL clearly separated slow stimuli (12.5-25 deg/s) from intermediate speed (50-100 deg/s). Areas AM and RL were specialized for speed discrimination at intermediate frequencies. Area LI showed an elevated discrimination power for slow speed pairs (3.1-100 deg/s). Area PM showed a similar yet less pronounced pattern comparing to LI. Moreover, the frequency-specific enhancement of speed discrimination performances in higher visual areas was not merely a consequence of individual areas’ frequency preferences, because the discriminability of cross-speed pairs remarkably outperformed the discriminability of iso-speed stimulus pairs in the same range of frequency space (Figure 5C; compare to Figure supplement 4C;). Altogether, these results highlight the area-specific enhancement of speed discriminability in higher visual areas, suggesting differential roles of higher visual areas in the visual motion processing.

### Asymmetric sampling of visual information channels in mouse visual areas

We next examined the representations of visual processing channels in mouse visual areas. In the previous sections we characterized independently how specific aspects of visual scenes are represented in individual visual areas. To identify visual processing channels and examine multidimensional tunings, we employed a non-supervised clustering approach (spectral clustering, see Method and Materials) to group neurons according to their responses to isotropic and anisotropic stimuli of different spatiotemporal frequencies. The clustering approach also allows to determine whether different visual feature subspaces are captured by distinct cell populations in the visual cortex, and how these functional cell types are represented across visual areas.

We found neuronal responses in visual areas can be divided into broad functional cell classes. To cluster responses, we repeatedly sampled randomly 2000 response vectors to give each area equal weight. Each response vector contained the amplitudes of responses to ISO and ANISO stimuli of all combinations of spatial and temporal frequencies (Figure 6A). To determine the number of clusters needed to capture the diversity of responses across visual areas, we computed the total within-cluster variability as a function of number of clusters. The resulting Scree plot showed an inflection point at 12 clusters (Figure supplement 5A), indicating that the population response patterns to visual stimuli in visual areas can be described by approximately 12 functional cell groups, representing a unique combination of visual features, such as distinct spatiotemporal tunings and differential responses to isotropic and anisotropic stimuli (Figure 6A–B; Figure supplement 6).

**Figure 6:**
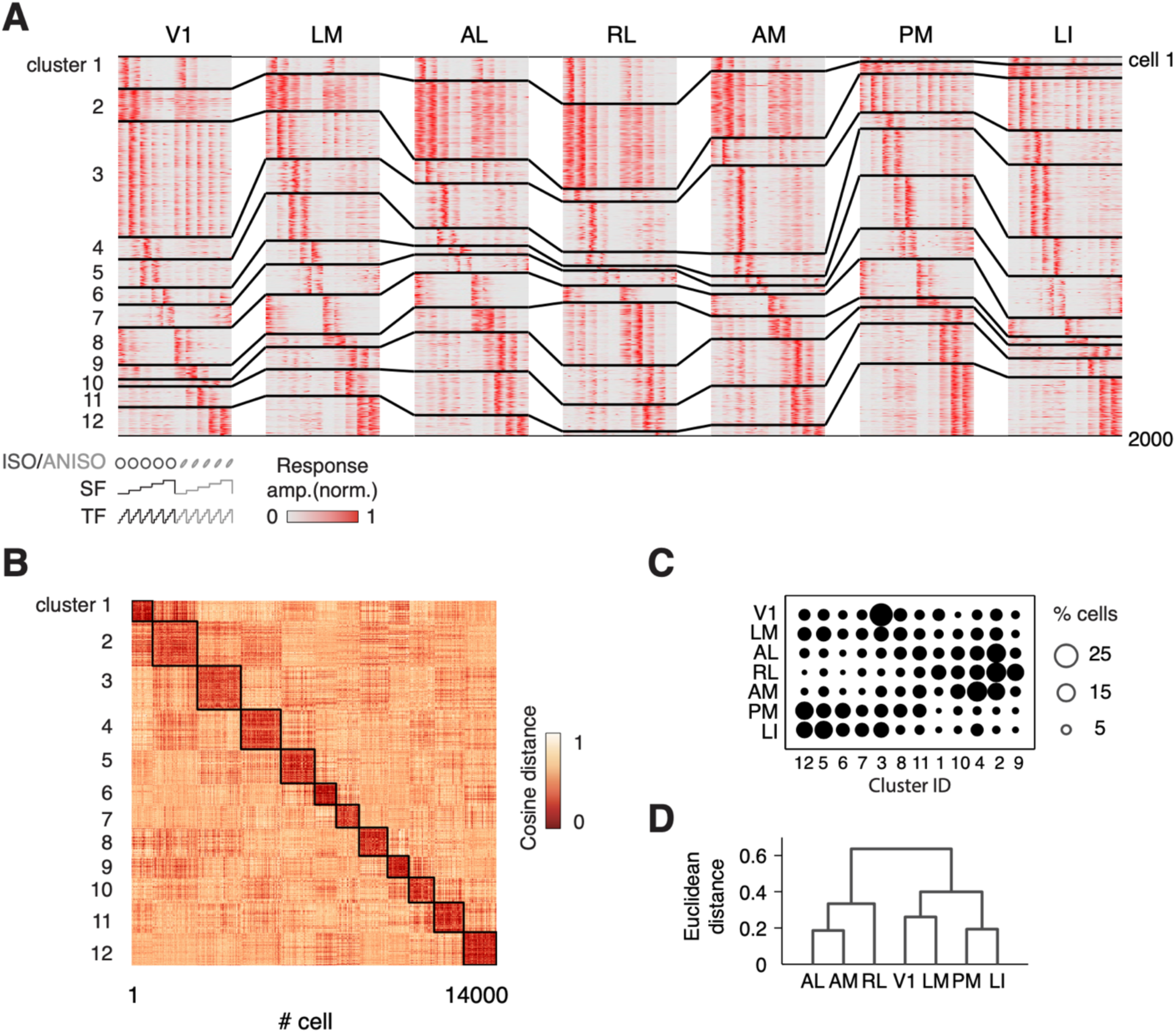
Asymmetric distribution of visual channels across cortical areas. (A) Matrices showing individual areas are characterized by distinct composition of functional cell types. Each matrix comprises the response profiles of 2000 randomly selected neurons. Each row represents a single cell’s peak-normalized responses to ISO and ANISO stimuli. Cells are clustered into 12 functional groups based on the similarity of response profiles using a spectral clustering approach (Materials and methods). The black lines delineate neighboring groups and connect the same groups across matrices. The stimulus parameters are annotated below the V1 matrix. (B) Heat map showing the cosine distances between the response profiles of the whole dataset. Black boxes delineate the 12 clusters. (C) Dot plot presenting the proportion of functional groups within and across areas. The groups are reordered to demonstrate the similarities and differences between areas. (D) The dissimilarity between areas are visualized as a Hierarchical tree based on the Euclidean distances estimated from the population composition across areas. The following figure supplements are available for figure 6: Figure supplement 5: Clustering. Figure supplement 6: Summary of functional groups.

We observed differential preferences for isotropic and anisotropic stimuli across identified cell groups (Figure supplement 6B), indicating distinct tuning for spatial elongation. To examine this, we further computed each group’s average tuning for stimulus anisotropy from the responses to elongated textures, which were not used for the clustering. The resulting tuning curves showed remarkable consistency with individual groups’ preferences for ISO versus ANISO stimuli (Figure supplement 6E), confirming independently that the dichotomy of selectivity for non-oriented versus oriented contrast is a distinguishing feature between functional groups.

Although each visual area encompasses cells belonging to each visual information channel, the proportion of cells belonging to each channel varied widely across visual areas (Figure 6C), such that each area was characterized by a distinct proportion of cells showing different isotropic and anisotropic preferences and spatiotemporal properties (Figure 6A). To quantify the similarities of visual representations in visual areas, we calculated the Euclidian distance of population compositions between pairs of areas, from which we computed a hierarchical tree of visual cortical areas (Figure 6D). This comparison revealed V1 to be most similar to area LM, showing relative uniform compositions of functional cell types. Areas PM and LI showed unique enrichment of some groups and sparsity of some other groups, positioning them in a different branch from V1 and LM. These patterns were reversed in anterior areas, showing a vast segregation from posterior areas (Figure 6C,D). These results indicate that mouse visual areas differentially sample from distinct spatiotemporal frequencies channels. These channels have simple stereotyped responses properties that capture a large fraction of the diversity seen across visual areas.

## Discussion

This study provides a detailed functional characterization of the visual features driving visual neuron responses and a comprehensive comparison of the cellular basis of representations across seven areas of the mouse visual cortex. Our findings reveal highly distributed visual information channels, representing distinct spatiotemporal and spatial orientation features in the visual cortex. Biased distribution of these processing channels across areas suggests differential role of individual areas in specialized visual computations. Considering the accumulating evidence of higher-order computations in the mouse higher visual cortex (Akrami et al., 2018; Burgess et al., 2016; Meijer et al., 2019; Morcos and Harvey, 2017), it is essential to understand how area-specific representations of visual features arise along the visual hierarchy, and how information from multiple modalities are integrated in the higher order cortex for complex computations, such as visual motion and shape processing. The results and implications of this study, provide a necessary basis for future studies investigating circuitry mechanisms for visual perception and behaviors.

### Comparison of spatiotemporal selectivity with previous studies

The preferred spatiotemporal frequencies observed in the current study are remarkably higher than in an earlier study in anesthetized mice (Marshel et al., 2011), but comparable to what has been reported in awake mice (Andermann et al., 2011). Moreover, our findings are not limited to a specific mouse line (Thy1-GCaMP6s mice), recordings we performed in another GCaMP6 reporter mouse line (TRE-GCaMP6s line G6s2) (Wekselblatt et al., 2016), which labels a broader population of pyramidal neurons, yielded similar results (data not shown). The lower ranges of preferred temporal frequencies observed in anesthetized mice (Marshel et al., 2011; Roth et al., 2012; Tohmi et al., 2014; Van Den Bergh et al., 2010) might result from suppressed thalamocortical synaptic transmission (Reinhold et al., 2015). Another factor that should be taken into consideration is the differential neuronal populations sampled in these two studies. In Marshel’s study (Marshel et al., 2011), excitatory and inhibitory neurons were ubiquitously labeled with the synthetic calcium indicator (Oregon Green Bapta-1 AM); whereas here we sampled from a subset of excitatory neurons in Thy1-GCaMP6s transgenic mice (Dana et al., 2014). As V1 interneurons prefer lower spatial frequencies than layer 2/3 excitatory neurons (Niell and Stryker, 2008), the population preference would shift towards lower frequencies with the inclusion of interneurons. However, it is unclear whether such a difference between excitatory and inhibitory neurons also exist in higher visual areas, and consequently leads to the lower preferred spatial frequencies as a population. Further comparisons of neuronal response properties in wakefulness and anesthesia (Adesnik, 2017; Durand et al., 2016; Greenberg et al., 2008; Haider et al., 2013) can explain the influence of different brain states on the neural representation and transformation among visual cortical areas. Nevertheless, studying neuronal physiology in the awake brain will be of great value for understanding neural computations for perception and behavior.

### Selectivity for stimulus anisotropy

We also found cortical neurons to be highly selective for stimulus anisotropy, suggesting diverse spatial integration properties. Many neurons show linear spatial summation (or ‘length summation’, (Schumer and Movshon, 1984)), such that their responses increase as the oriented textures elongate. Meanwhile, many neurons prefer short stimuli, reminiscent of ‘end-stopping’ cells, which preferentially respond to stimuli of certain lengths (Hubel and Wiesel, 1965; Gilbert, 1977). This length tuning property may be attributed to ‘surround suppression’, where one receptive field is inhibited by the stimulation at the surround. Diverse length tuning curves may result from differential ratios of the excitation on classical receptive fields and the inhibitory effect of the receptive field surround (Adesnik, 2017; Adesnik et al., 2012; Vaiceliunaite et al., 2013).

We found diverse spatial integration properties distributed across the cortical areas with area-specific biases, which might participate in distinct neural computations. V1 and LM show relatively strong biases for non-oriented stimuli, suggesting strong surround suppression that is also observed in the primate (El-Shamayleh et al., 2013; Hubel and Livingstone, 1987; Shushruth et al., 2009), the cat (DeAngelis et al., 1994) and the mouse (Adesnik, 2017; Adesnik et al., 2012; Murgas et al., 2020; Nienborg et al., 2013; Vaiceliunaite et al., 2013; Van Den Bergh et al., 2010). Surround suppression is involved in a number of neural computations, including shape processing. In the primate, ‘end-stopping’ behavior increases along the ventral processing stream and is suggested to benefit the coding for curvatures (Ponce et al., 2017). Recent studies also revealed the lateral areas in the rat also show less sharpness of the orientation tuning and insensitivity to the phase of the drifting gratings, which might contribute to the invariant representations of visual shapes of different scales and shapes in these lateral areas (Matteucci et al., 2019; Tafazoli et al., 2017; Vermaercke et al., 2014). These observations agree to the strong preferences for non-oriented stimuli in the lateral area LI observed in this study. Future studies characterizing the role of nonlinear spatial integration properties in building invariant representations of visual shapes will benefit the understanding of computational mechanisms of object processing.

### Functional organization of mouse higher visual areas

Our findings revealed a clear asymmetry in representation of the spatiotemporal features in the mouse visual cortex which may indicate a functional organization that is adapted to encode ethologically relevant visual information. We uncovered a graded representation of fast to slow motion along the anterior-posterior axis in the mouse higher visual cortex. Anterior areas, especially areas RL and A, mainly represent the lower visual field (Zhuang et al., 2017). The preferences for large-scale, fast motion of these areas might benefit the encoding for fast optic flows near the ground during animal’s exploratory behaviors. Supporting this theory, a recent study found area RL is specialized for binocular disparities corresponding to nearby visual stimuli (La Chioma et al., 2019) and might contain a visuo-tactile map of near space (Olcese et al., 2013). In contrast, area PM mainly represents the visual periphery, which is dominated by strong but relatively slow visual flows during animals’ navigation. In an additional set of experiment (data not shown), using isotropic noise stimuli with global motions, we found PM neurons, with the similar preference for slow speed, respond more robustly to coherent, global motion than to incoherent, local motion, suggesting the specialization for encoding slow optic flows in the visual periphery during navigation. Area PM might relay information about optic flows via the dense, direct efferent projections to the retrosplenial cortex (Wang et al., 2012), which preferentially responds to slow motion (Murakami et al., 2015) and is involved in spatial navigation (Mao et al., 2017). Moreover, area-specific encodings of behavioral variables and the distinct projections of dorsal areas suggest their differential involvements in visuomotor behaviors (Minderer et al., 2019; Wang et al., 2012; Wang and Burkhalter, 2013). Altogether, these data suggest the dorsal higher visual areas of mouse are specialized for ethological encoding of visual motion, reminiscent of the dorsal streams in the primate visual system (Van Essen and Maunsell, 1983).

The ventral processing stream has been found a conserved organization across many species, including primates, cats and rats (Matteucci et al., 2019; Tafazoli et al., 2017; Van Essen and Maunsell, 1983; Vermaercke et al., 2014). Here we found functional evidence suggesting it also exists in the mouse lateral cortex. Lateral area LI shows high spatial acuity and ‘end-stopping’ properties. These properties are suitable for encoding spatial details and curvatures, which are critical for object recognition (Lu et al., 2018; Ponce et al., 2017). However, in the mouse visual system the presence of higher-order functions, such as transformation-tolerant representation of visual objects (Tafazoli et al., 2017; Vermaercke et al., 2014), has not yet been tested. Other lateral areas, including postrhinal area (POR) and posterior area, were hinted to have similar preferences for low temporal and high spatial frequencies (Murakami et al., 2017), consistent with the anterior-posterior cortical gradient for fast-to-slow motion information. Recent studies suggest POR as a hub region, associating visual objects with behaviorally relevant variables, such as reward values and object location (Burgess et al., 2016; Furtak et al., 2012). How do these properties coordinate with other visual properties, such as selectivity for moving objects (Beltramo and Scanziani, 2019; Bennett et al., 2019) and for slow stimuli (Murakami et al., 2017), and the behavioral implications of such a functional specialization remain to be determined. Strong anatomical connections of these lateral areas to temporal and parahippocampal regions (Wang et al., 2012), supports the idea that mouse lateral visual cortex might be the involved in shape processing, although species-specific differences might also exist.

Based on the feedforward and feedback characteristics of the intracortical connectivity, (Felleman and Van Essen, 1991), area LM is generally placed at the first stage of higher-order visual processing (D’Souza et al., 2016; Harris et al., 2019). It receives strong feedforward inputs from V1 (Glickfeld et al., 2013; Wang and Burkhalter, 2007) and in return provides strong feedback projections, which might facilitate predictive coding of sensory information (Keller et al., 2020; Nguyen et al., 2018). Area LM has been proposed to be the gateway for the ventral stream (AL as the gateway for the dorsal stream), given its relatively denser projections to the ventral areas (Wang et al., 2012, 2011) and population responses similar to the ventral areas (wide field imaging in Murakami et al., 2017; Smith et al., 2017). However our data show that LM contains highly diverse neurons that exhibit both dorsal and ventral properties, resembling the primate V2 (Van Den Bergh et al., 2010). In addition, LM neurons send strong projections to both dorsal and ventral areas (Wang et al., 2012, 2011), routing target-specific information (Glickfeld et al., 2013). These functional and anatomical characteristics argue for LM’s role as the divided gateway of dorsal and ventral streams.

The functional specialization of visual cortical areas must depend on specific intracortical connectivity. Previous anatomical studies revealed a nearly ‘all-to-all’ diagram of intracortical connections in the primate and rodent visual cortex (Gamanut et al., 2018; Wang et al., 2012), suggesting intensive crosstalk between functionally specialized areas. The information transmitted through the connections, however, remain unclear. Individual projection neurons either broadcast the same information across multiple areas, or selectively target a single area (Han et al., 2018). The connectivity between higher visual areas can be functionally target-specific, in the similar manner as the projections from lower to higher cortical areas (Glickfeld et al., 2013; Kim et al., 2018). Therefore, neurons sharing similar response properties might interconnect via long-range projections and form cortical subnetworks that spread the common information. Here we uncovered the widespread distribution of shared response types across the visual cortex, suggesting multiple distributed cortical subnetworks. The similarity of response type compositions between areas might be predictive of the strength of anatomical connectivity between visual cortical areas (Gamanut et al., 2018; Wang et al., 2012). Future studies combining *in vivo* cellular physiology along with circuit mapping and manipulation should provide important insights to the organization rules of connectivity and information flow between visual cortical areas.

## Materials and Methods

### Animals and Surgery

All experiments were approved by the Animal Ethics Committee of KU Leuven. Standard craniotomy surgeries were performed to gain optical access to the visual cortex (Goldey and Andermann, 2014). C57BL/6J-Tg (Thy1-GCaMP6s)GP4.12Dkim/J mice (Dana et al. 2014; 5 male and 5 female) between 2 and 3-month-old were anesthetized with isoflurane (2.5%–3% induction, 1%–1.25% surgery). A custom-made titanium frame was mounted to the skull, and a craniotomy was made to expose the left visual cortex. The 5mm-wide craniotomy was covered by a set of cover glasses. Buprenex and Cefazolin were administered postoperatively when the animal recovered from anesthesia after surgery (2 mg/kg and 5 mg/kg respectively; every 12 hours for 2 days).

### Behavior monitoring

All mice were habituated at least 3 days before data acquisition. During imaging experiments, mice were head restrained on a treadmill, passively viewing the display directed to the right eye. We recorded animals’ right eye movement using a macro lens (Zoom 7000, Navitar) mounted to a GigE camera (Prosilica GC660, Allied Vision Technologies), we tracked the running speed and monitored the animals’ general behavior (MAKO G-030b, Allied Vision Technologies). Mice showed no consistent behavioral or gaze responses to the visual stimulation.

### Visual Stimulation

Visual stimuli were generated in Matlab (The Mathworks) and delivered with customized software written in python. Stimuli were spherically corrected and displayed on a gamma-corrected LCD display (22’’, Samsung 2233RZ). The screen was oriented parallel to the right eye of the animal with 18 cm in distance (covering 80 deg in elevation by 105 deg in azimuth).

Spectral noise stimuli (Figure1D, 2A -- Figure supplement 1D) were created by applying a set of parametrized filters to random pink-noise fields. Narrow bandpass filters (bandwidth: 1 octave) for spatial and temporal frequencies determine the scale and flickering frequencies of the textures. The orientation bandwidth determines the length/width ratio of spatial patterns, which we refer to as stimulus anisotropy. Isotropic noise stimuli, which are spectrally band-passed noise with no angular filter, resemble clouds of moving dots (Figure supplement 1D upper row; ISO stimuli). Narrowing the orientation bandwidth leads to elongation of textures, yielding higher stimulus anisotropy (Figure supplement 1D). While isotropic textures contain no global motion, oriented textures were designed to slowly rotate (45 deg per second, clockwise) to activate neurons tuned to different orientations.

This study uses three stimulus sets generated by combining the above stimulus dimensions. For the spatiotemporal frequency assay, we used ISO and ANISO stimulus sets, which contained non-oriented and highly elongated textures (orientation bandwidth: 5 deg) respectively (Figure 1D). Both stimulus sets comprised 30 combinations of spatial frequencies (0.02, 0.04, 0.08, 0.16, cpd) and temporal frequencies (0.5, 1, 2, 4,8,16 Hz). For the stimulus anisotropy assay, the stimulus set contains the combinations of 4 spatial frequencies (0.04, 0.08, 0.16, 0.32 cpd) and 4 orientation bandwidths (infinite, 40, 10, 5 deg FWHM), with a fixed temporal frequency at 2 Hz (Figure 2A,B). Each stimulus condition was presented for 4 second, intertwined with 4-second equiluminant gray screen. In each of the four pseudorandomized trials, a different seed was used to generate a unique set of random pink noise, with varying phases yet constant frequency spectra across trials.

### Two-photon Calcium Imaging

A customized two-photon microscopy (Neurolabware) was used. The calcium indicator GCaMP6s was excited by a Ti:Sapphire excitation laser (MaiTai DeepSee, Spectra-Physics) operating at 920 nm. The green fluorescence of GCaMP6s was collected with a photomultiplier tube (PMT, Hamamatsu) through a bandpass filter (510/84 nm, Semrock). Images (720×512 pixel per frame) were collect at 31 Hz with a 16x objective (Nikon). Volume imaging was achieved using a focus tunable lens (EL-10-30-TC, Optotune; staircase mode). We recorded activity from neurons typically between 100 to 300 µm deep below pia.

### Widefield Calcium Imaging

Widefield fluorescent images were acquired with a 2x objective (NA = 0.055, Mitutoyo, Edmund Optics). Blue LED light (479nm, ThorLabs) was used for illumination passing an excitation filter (469/35 nm Thorlabs) and a dichroic filter centered at 498 nm. The green fluorescence was collected with an EMCCD camera (EM-C^2^, QImaging) via a bandpass filter (525/39 nm filter, Thorlabs). The image acquisition was controlled with customized software written in python.

### Retinotopic mapping

To identify visual cortical areas *in vivo*, we performed retinotopic mapping with 1-photon wide field imaging using dynamic noise stimuli partially visible through a circling round aperture (“circling patch”) or a translating rectangular aperture (“traveling bar”). The circling patch stimuli is a small patch of contrast (20 deg in diameter) circling along an elliptic trajectory on the display (azimuth: -40 to 40 deg; elevation: -30 to 30 deg; 20 sec per cycle with 20 repetitions; Figure supplement 1 A). The traveling bar stimuli comprised a narrow bar (13 deg wide) sweeping across the screen in 4 cardinal directions (10 sec per sweep with 20 repetitions). An isotropic noise background (0.08 cpd, 2 Hz) was embedded in the patch or bars. The phase maps were generated by calculating the phase/position of the stimuli evoking maximal neural responses for each image pixel. Each area has a complete representation of the elliptic trajectory of the circling patch, resulting pinwheel-like retinotopic maps (Figure supplement 1B). The traveling bar stimuli generated azimuth and altitude phase maps, which were used to generate sign maps with the methods described in the previous study (Figure supplement 1C) (Juavinett et al., 2017). Areas are identified according to the common delineation (Wang and Burkhalter, 2007; Zhuang et al., 2017). The boarders between areas were further refined with 2-photon imaging of cellular responses to retinotopic stimuli to assign neurons into different areas.

## Data Analysis

All subsequent data analysis was performed in MATLAB (The Mathworks, Natick, MA).

### Selection for visual cells

For the cellular imaging, raw images were reconstructed and corrected for brain motion artefacts using custom MATLAB routines (Bonin et al., 2011). Regions of interest (ROI) were selected at somatic regions showing significant calcium activities during visual stimulation. Cellular fluorescence time courses were generated by averaging all pixels in a ROI, followed by subtracting the neuropil signals in the surround shell. Responses were defined as the averaged ΔF/F during the stimulus epoch, where ΔF is the change in the fluorescent signal and F is the baseline fluorescence (1s before stimulation). Stimulus-evoked responses were identified when median cross-trial response curves showed response magnitude higher than 3x standard deviation of the baseline activity for over one second. A cell was classified as responsive if it shows stimulus-evoked responses to at least one stimulus condition. Response reliability r was measured as the 75 percentiles of the correlation coefficients between stimulus-evoked fluorescence traces across trials. Neurons above the threshold of 0.3 were deemed reliable.

### Tuning curves

The anisotropy preference index (API) is expressed as ^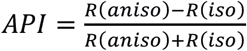^, where R(aniso) and R(iso) are individual neurons’ peak response amplitudes to ANISO and ISO stimuli respectively. For the spatial and temporal frequency assay, responses were fit to two-dimensional elliptical Gaussian models (Priebe et al. 2003; Andermann et al. 2011):

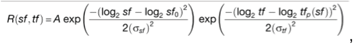

where log_2_ *tf*_*p*_ (*sf*) =ξ (log_2_ *sf* − log_2_, *sf*_0_) + log_2_ *tf*_0_ and A is the peak response amplitude, sf_0_ and tf_0_ are the preferred spatial and temporal frequencies, and *σ*_*sf*_ and *σ*_*tf*_ are the spatial and temporal frequency tuning widths. The dependence of temporal frequency tunings on the spatial frequency is captured by a power-law exponent ξ.

Neurons are categorized as lowpass, bandpass and highpass based on whether the responses to the minimal and maximal tested frequencies are greater than half maximal responses. Lowpass cells tend to respond to frequencies lower than the tested spectrum; while highpass cells are likely to respond to higher frequencies. Bandpass neurons respond mainly within the tested spectrum.

Estimates of cutoffs for the spatial and temporal frequency were measured at cross-sections at *R(sf, tf*_*0*_) and *R(sf*_*0*_, *tf)*, respectively. We estimated low cutoffs from bandpass and highpass cells, high cutoffs from bandpass and lowpass cells, and bandwidths from bandpass cells.

### Population coding analysis

To examine the information encoded in individual visual cortical areas, we decoded stimulus categories from neuronal population activities using linear classifiers (Hung et al., 2005; Vermaercke et al., 2014). Support vector machines were trained and tested in pair-wise classification for all possible pairs (30 stimulus conditions result in 435 unique pairs). To compare the decoding performance with the result of cell-by-cell speed tuning analysis, we focused on the neurons responded to ISO stimuli. Visual responses across four trials were split into training and testing groups (half-half) and the performance was measured as the proportion of correct classification decisions to the testing groups (standard cross validation). The decoding analysis were repeated for 100 times with random sampling. To test the scaling of decoding performance as a function of population size, we measured the decoding performances as a function of numbers of sampled neurons (logarithmic increase from 1 to 1000; without replacement) per area. To compare the discriminability for specific stimulus pairs, we measured the number of neurons required to reach the classification accuracy at 90% by interpolate the growth curves of performances as a function of population size. To assess the differences in the overall and feature specific decoding performance between visual areas, we compared the decoding performances between iso-speed stimulus pairs and between cross-speed pairs (Figure 6C – Figure supplement 4C). Iso-speed pairs were neighboring textures lying along the iso-speed lines. Cross-speed pairs are neighboring textures lying orthogonally to iso-speed lines and have a 4-fold difference in speed.

### Clustering

To assess whether cortical neurons form functional cell types, we used a spectral clustering approach. We randomly selected 2000 neurons with replacement from individual areas to avoid bias. Each neuron is expressed as a vector of responses to ISO and ANISO stimuli of all combinations of spatial and temporal frequencies (response amplitude normalized to range from 0 to 1). The response matrix of all selected neurons was analyzed using a spectral clustering algorithm (Ingo, 2020). To determine whether there are discrete functional types, we plotted the overall variances (within-cluster sum-of-square variances) as a function of number of clusters. The resulting Scree plot showed an inflection point at 12 clusters (Figure supplement 5A), suggesting 12 broad functional cell types. We measured the similarity between neurons within individual clusters and across clusters, based on the cosine distance between response vectors, and found high similarities within clusters and high differences between clusters. These clusters showed distinct response properties and differential distributions across areas (Figure supplement 6). To quantify the similarity/differences between visual areas in terms of the proportions of functional cell types, we calculated the average Euclidean distances between areas and expressed them in a dendrogram (Figure 6D). We ran the clustering algorithm, with a constrain of 12 clusters, with random sampled pools for 100 times. The resulted dendrograms were highly consistent (Figure supplement 5C).

## Acknowledgements

We are grateful to members of the Bonin and Farrow laboratories for feedback, discussions and comments on the manuscript, Joao Couto for establishing the imaging setup and support with image processing, Karl Farrow and Mark Andermann for comments on an earlier version of the manuscript, and Asli Ayaz for editing and valuable suggestions for the manuscript writing. This work was supported by Neuro-Electronics Research Flanders (VB), Research Foundation Flanders (FWO) fellowship 12E4314N (BV), grant G0D0516N (VB), KU Leuven Research Council grant C14/16/048 (VB) and a personal fellowship award from Chinese Scholarship Council (XH).

## Author Contributions

Conceptualization & Methodology, X.H., B.V. and V.B.; Investigation, X.H. and B.V.; Data Analysis, X.H., B.V, and V.B. ; Writing – Review & Editing, X.H., B.V. and V.B.; Funding Acquisition, V.B.; Resources, V.B.; Supervision, V.B.

