## Supplementary Figures for "Cellular organization of visual information processing channels in the mouse visual cortex"

### Supplementary figure legends

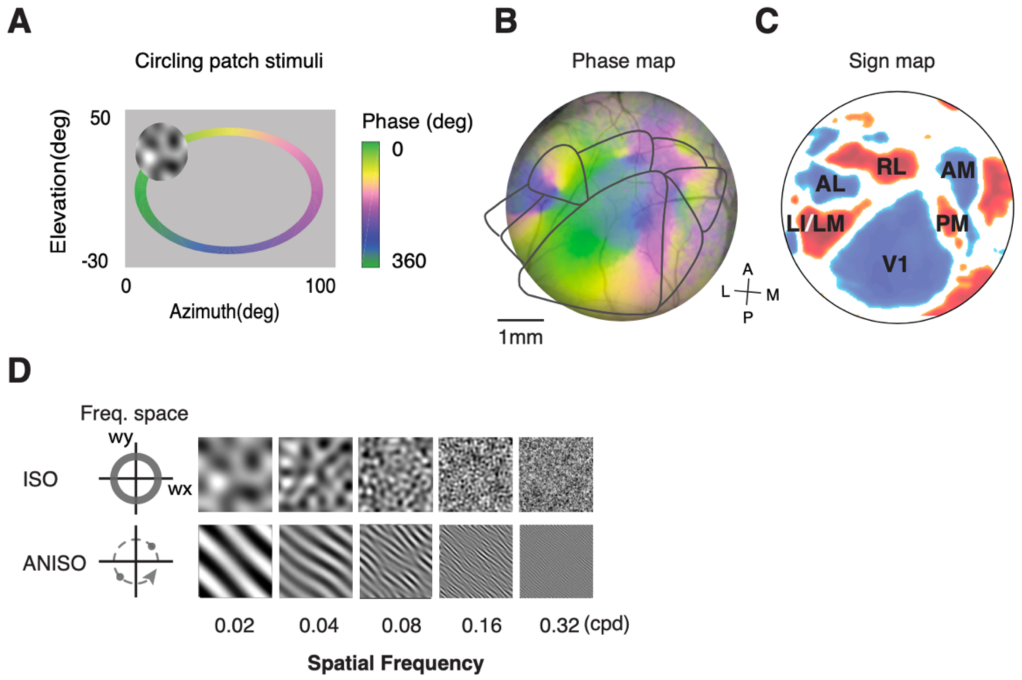


**Figure supplement 1: Retinotopic mapping and spectral noise stimuli.** (A) Scheme of visual stimulation. A patch of stimuli with a spectral noise background (0.08 cpd, 2Hz) continuously circles on the display along an elliptic trajectory. The phase of the trajectory is shown in color. (B) Phase map showing cortical regions responding to different phases of the trajectory. Each area has a representation of the full trajectory, resulting in a 'pinwheel' retinotopic map. (C) Sign map showing visual cortical areas (blue and red patches). (D) Spectral noise stimuli with varying spatial frequencies.

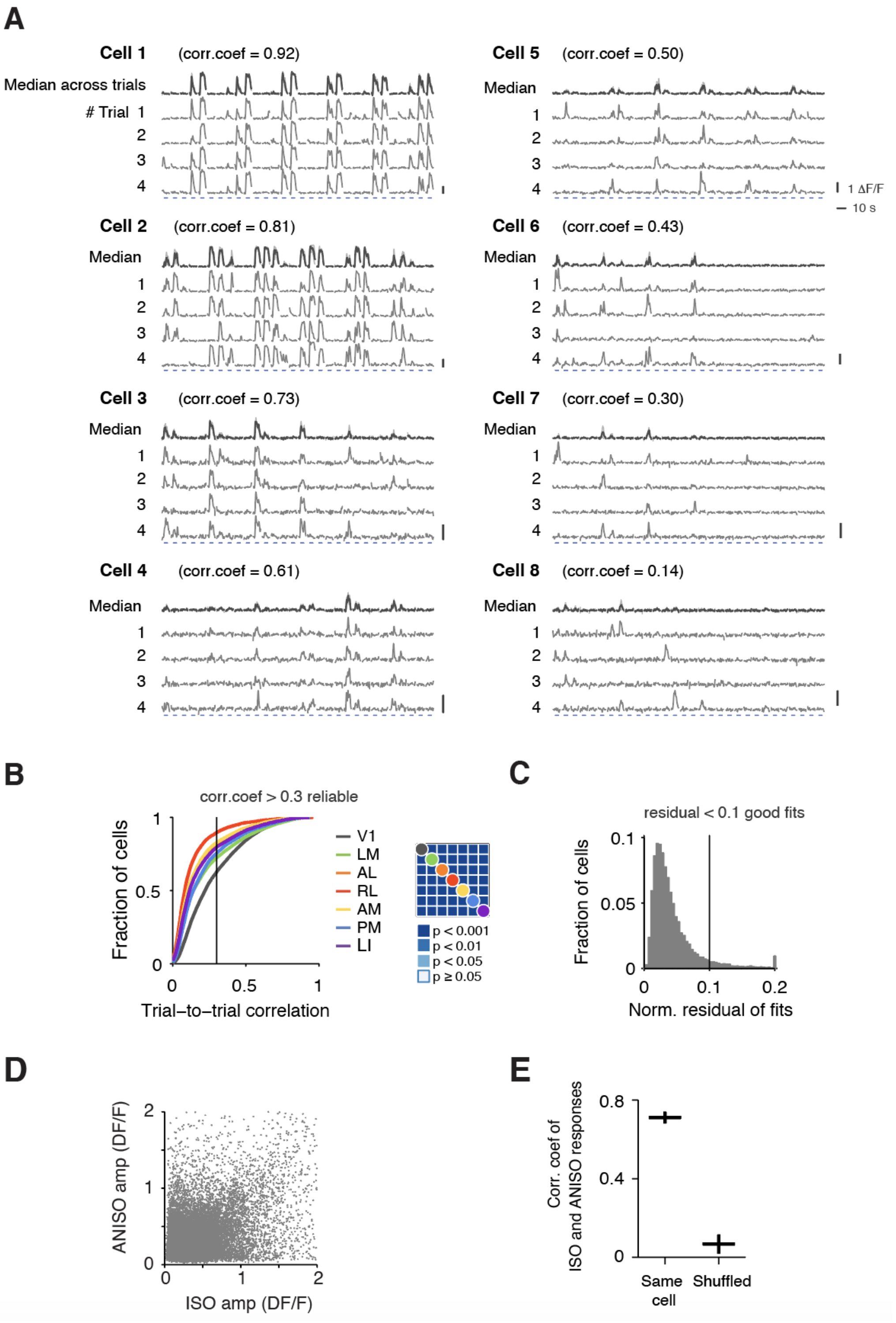


**Figure supplement 2: Population statistics of response to ISO and ANISO stimuli.** (A) Example cells showing responses across trials with different responsiveness and variabilities. The top trace of each cell shows the median responses across trials. The median absolute deviations across trials are shown as light gray shadow. Blue bars indicate 4s stimulus epochs. Gray scale bars indicate 1 $\Delta F/F$ and 10 second. The $75^{th}$ percentile trial-to-trial correlation coefficients are noted above the response traces. (B) Cumulative distributions of the $75^{th}$ trial-to-trial correlation coefficient of the populations across areas. Cells with a value above 0.3 are considered reliable. V1 shows higher reliability than higher visual areas. All pairs are significantly distinct (KS test with Bonferroni correction). (C) Histogram showing the distribution of the normalized residual of spatiotemporal tuning fits. Fits below the threshold at 0.1 were considered good fits and included in further analysis. (D) Scatter plot showing high diversity of individual neurons' response amplitudes to ISO and ANISO stimuli. (E) Mean correlation coefficients of the responses to ISO and ANISO stimuli of cells responding to both stimuli versus shuffled pairs. Mean ± sem.

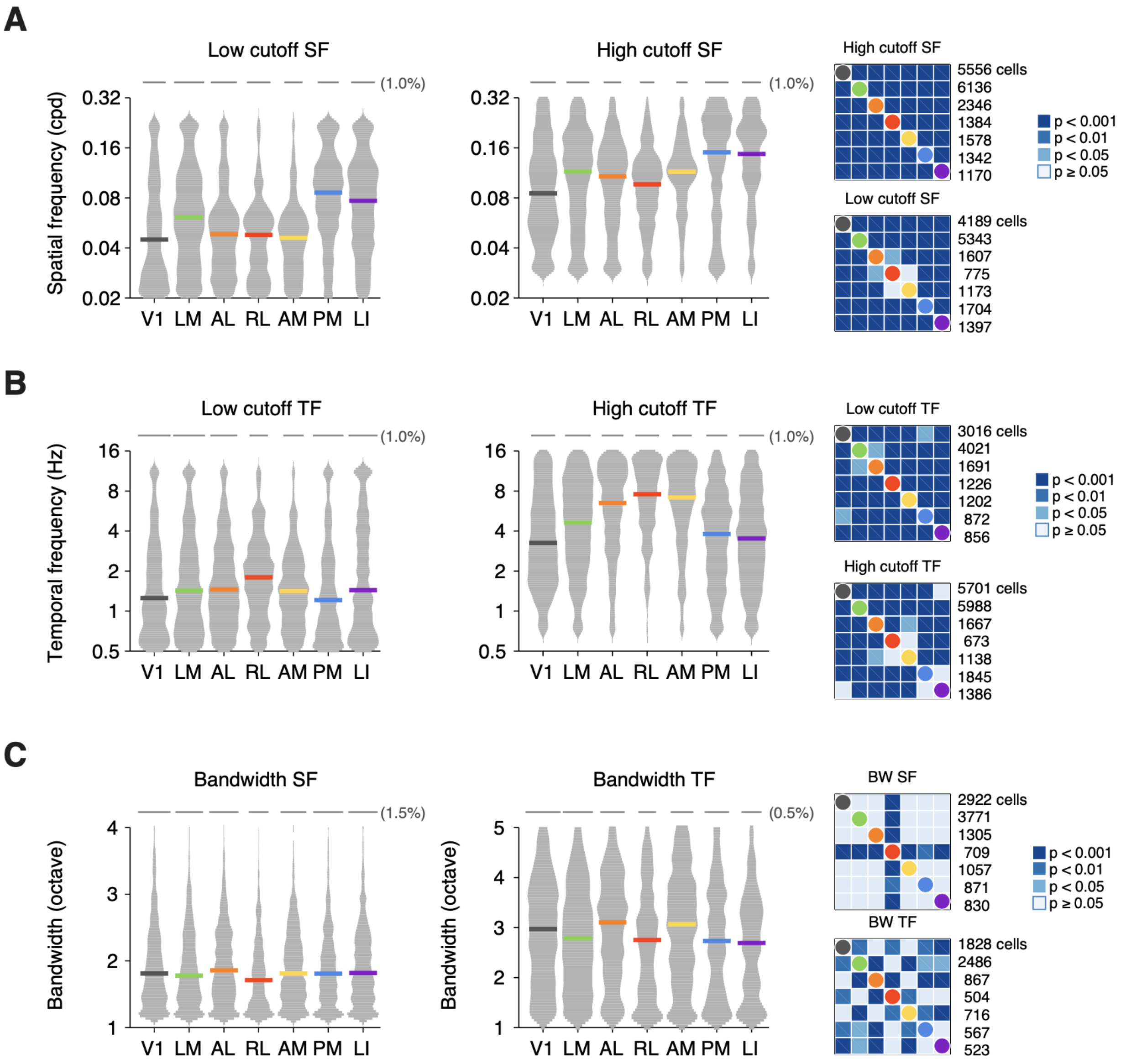


**Figure supplement 3: Spatiotemporal tuning parameters**. (A) Distribution of low and high cutoffs of spatial frequency across areas. Distributions are smoothed with a Gaussian kernel (bandwidth=0.1). Color bars indicate median values. Gray scale bars indicate 1% of cells for each distribution. Right panels: Statistic significances of pairwise comparisons. KS test with Bonferroni correction. (B) Same as (A), for temporal frequency. (C) Distribution of spatial and temporal tuning bandwidths.

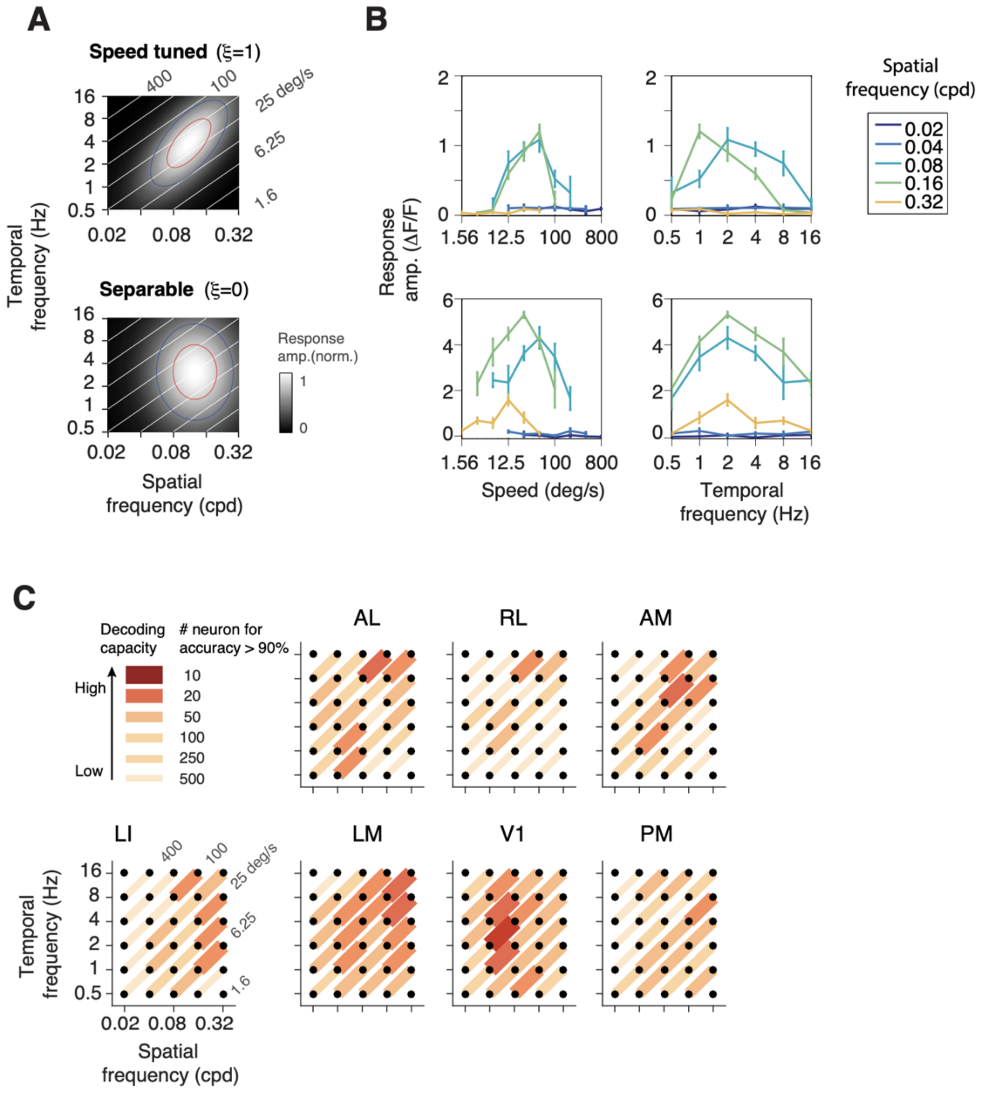


**Figure supplement 4: Speed tuning analysis.** (A) Example neurons that are speed tuned or untuned. The speed tuning index ξ describes the interdependency between temporal frequency and spatial frequencies. If ξ close to 1, the cell is tuned for speed (upper left); if ξ close to 0, the cell has separable tuning for spatial and temporal frequencies (red and blue contours show 80 and 50% peak amplitude; white lines indicate iso-speed lines). (B) Speed-tune cells have similar tuning curves for speed at different spatial frequencies (upper left), while the temporal frequency tuning curves change across spatial frequencies (upper right). Neurons with separable spatiotemporal tuning have similar tuning curves for temporal frequencies but not speed at different spatial frequencies (lower panels). (D) Discrimination tasks were performed between iso-speed pairs of isotropic stimuli. The discriminability is estimated as the number of cells required to reach 90% classification accuracy (red lines).

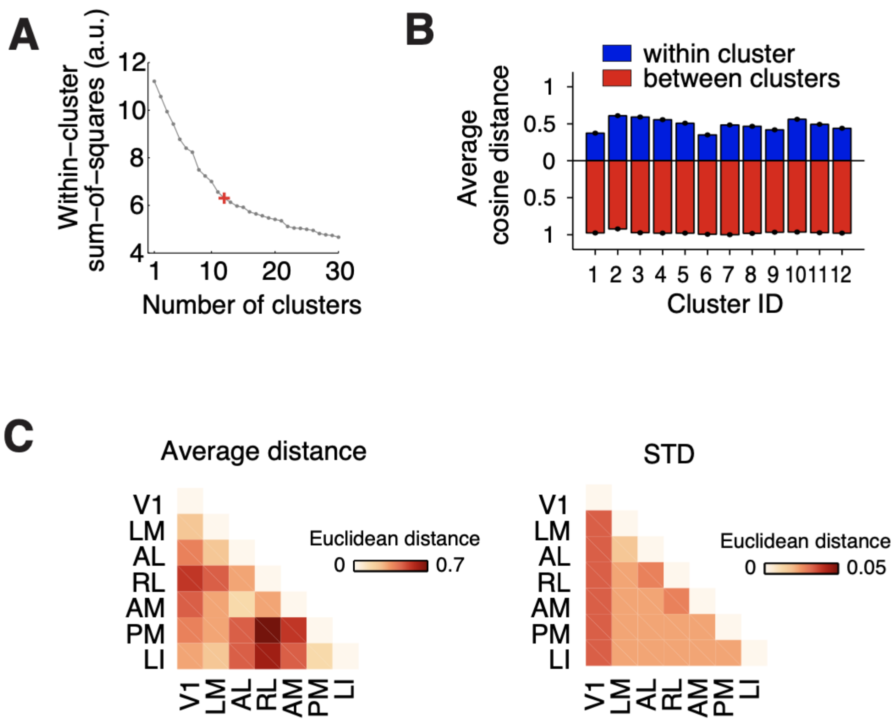


**Figure supplement 5: Clustering.** (A) Scree plot of the within-cluster sum-of-square as a function of number of clusters, showing an inflection point at 12 clusters. (B) Bar graph showing average cosine distances between pairs of cells within clusters (blue) and between clusters (red). (C) The left matrix shows the mean Euclidean distance of cell composition between areas averaged across 100 independent repetitions. The right matrix shows corresponding standard deviations.

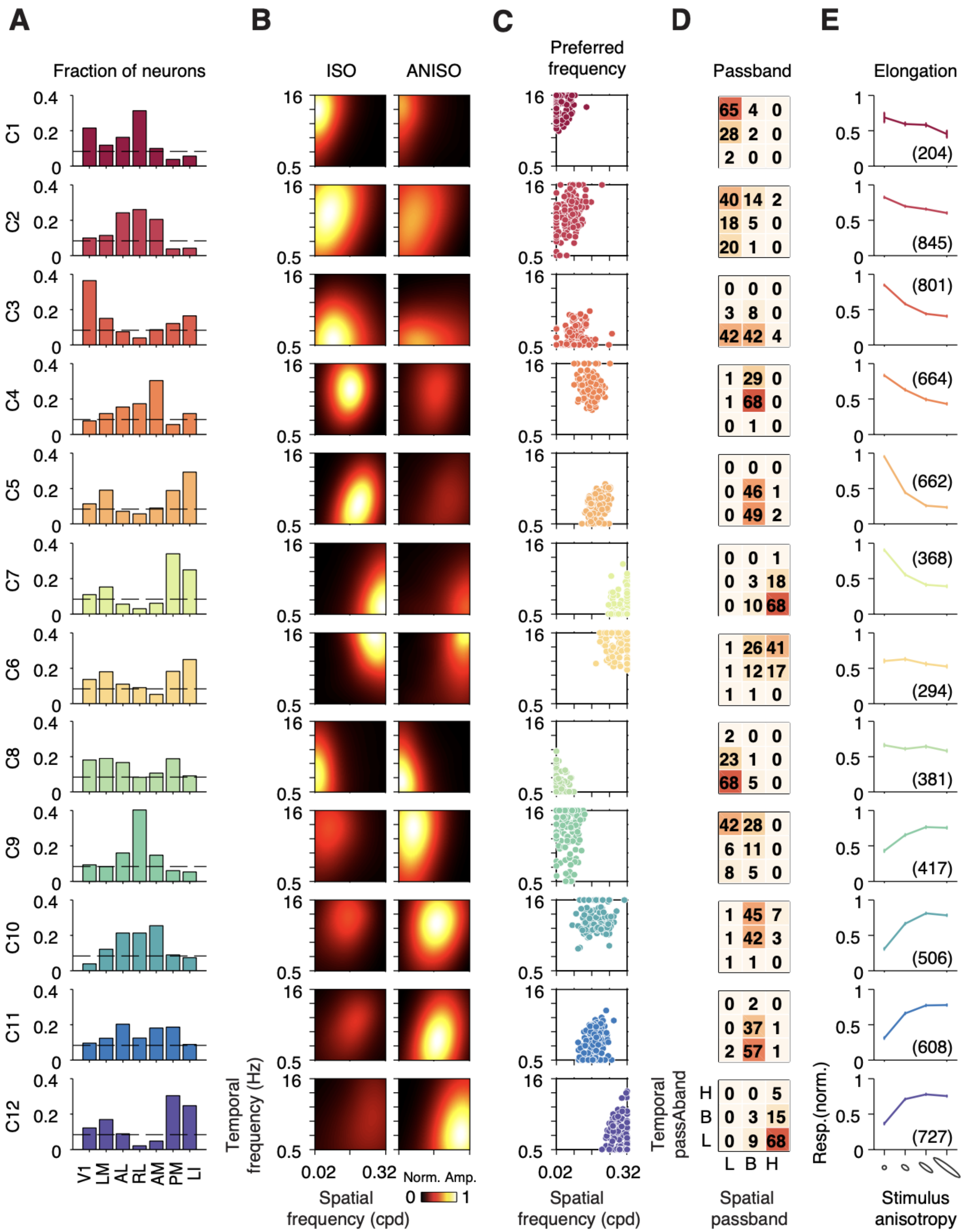


**Figure supplement 6: Summary of functional groups.** (A) Distribution of individual clusters across areas. The dash line shows the fraction of one cluster if all 12 clusters are equally represented (1/12). (B) Heat maps showing the average of peak-normalized fits of the responses to ISO and ANISO stimuli. (C) Scatter plots showing the joint distribution of preferred frequencies. (D) Passband properties for the spatiotemporal frequency. (E) Average tuning curves for stimulus anisotropy, showing consistency with the preferences for ISO or ANISO stimuli.
